# Uncured PDMS Inhibits Myosin In Vitro Motility in a Microfluidic Flow Cell

**DOI:** 10.1101/418327

**Authors:** Yihua Wang, Thomas P. Burghardt

**Affiliations:** Department of Biochemistry and Molecular Biology, Mayo Clinic Rochester Rochester, MN 55905; Department of Physiology and Biomedical Engineering, Mayo Clinic Rochester Rochester, MN 55905

**Keywords:** Myosin in vitro motility, Nanoliter volume motility assay, Polydimethylsiloxane/glass hybrid microfluidic, Unpolymerized PDMS inhibits myosin, Curing recovers myosin motility

## Abstract

The myosin motor powers cardiac contraction and is frequently implicated in hereditary heart disease by its mutation. Principal motor function characteristics include myosin unitary step size, duty cycle, and force-velocity relationship for translating actin under load. These characteristics are sometimes measured in vitro with a motility assay detecting fluorescent labeled actin filament gliding velocity over a planar array of surface immobilized myosin. Assay miniaturization in a polydimethylsiloxane/glass (PDMS/glass) hybrid microfluidic flow channel is an essential component to a small sample volume assay applicable to costly protein samples however the PDMS substrate dramatically inhibits myosin motility. Myosin in vitro motility in a PDMS/glass hybrid microfluidic flow cell was tested under a variety of conditions to identify and mitigate the effect of PDMS on myosin. Substantial contamination by the monomeric species in polymerized PDMS flow cells is shown to be the cause of myosin motility inhibition. Normal myosin motility recovers by either extended cell aging (∼20 days) to allow more complete polymerization or by direct chemical extraction of the free monomers from the polymer substrate. PDMS flow cell aging is the low cost alternative compatible with the other PDMS and glass modifications needed for in vitro myosin motility assaying.

## INTRODUCTION

The sliding actin in vitro motility assay measures myosin translocation of actin [1]. It is widely used to characterize cardiac myosin force-velocity [2; 3] and applicable for comparing normal to inheritable disease compromised cardiac function [4]. This assay in many configurations has uniquely low demand for the protein reagents allowing novel applications to contractile proteins extracted from human muscle biopsy samples [5] and to expensive expressed human myosins [6; 7]. The glass flow cell in the conventional in vitro motility assay has 10-50 µl volumes [1; 8]. Reducing assay volume from microliter to nanoliter scale involves assay adaptation to a microfluidic.

As one of the most popular materials used in the fabrication of micro- or nanoscale devices, poly(dimethylsiloxane) (PDMS) has many advantages, such as simple manufacturing methods, optical transparency, flexibility, gas permeability, ease of bonding to glass, and low cost [9; 10; 11]. Pristine PDMS is known to have poor compatibility with motor proteins [12; 13; 14]. We found that in a PDMS/glass hybrid microfluidic flow cell using pristine PDMS, where in vitro motility is detected with the myosin immobilized at the nitrocellulose (NC) coated glass surface, most actin filaments do not move. Just the presence of the PDMS in the cell inhibited motility. Neutralizing this effect is essential for its application in a microfluidic based assay.

Many PDMS surface modification methods have been developed such as plasma treatment, layer by layer (LBL) deposition, surfactant treatment, protein adsorption, silanization, chemical vapor deposition (CVD), graft polymer coating and hydrosilylation-based PDMS surface modification [15; 16; 17; 18]. We examined in vitro motility of skeletal heavy meromyosin (sHMM) in PDMS/glass hybrid chambers treated with several PDMS surface modification methods. We selected methods that were simple and convenient for routine practices in biochemistry and biomedical laboratories and had minimal requirements for specialized instruments. The methods include coating PDMS surface with NC, air plasma treatment, layer by layer deposition with poly(vinyl alcohol) (PVA)/glycerol [19], solution-phase oxidation [20], surfactant treatment using n-Dodecyl-β-D-maltoside (DDM) [21], chemical extraction of unpolymerized substrate, and 20 day aging treatment. The latter two treatments are not surface modifications but methods to remove or reduce unpolymerized substrate.

We found that using 20 day aged PDMS substrate to construct flow channels neutralized the inhibition of motility by PDMS and was the overall best approach. Using this method, we developed a PDMS/glass hybrid motility assay with a flow channel volume of 0.03 µl. Its performance compared favorably with the conventional glass motility flow chamber with a volume of 10-50 µl. In addition, results suggested that the contamination from the leaching of uncured PDMS oligomers into the sample media is the most significant inhibitor of in vitro myosin motility.

## MATERIALS AND METHODS

### Materials

Sylgard 184 PDMS base and curing agent were from Dow Corning (Midland, MI). Quantum dot 585 streptavidin conjugate (Qdot), rhodamine-phalloidin (Rh-ph), biotin-XX-phalloidin, and phalloidin were obtained from Life Technologies (Grand Island, NY). Glucose oxidase was purchased from MP Biomedicals (Santa Ana, CA). Biotin free bovine serum albumin (BSA, cat # A3059) and catalase were from Sigma-Aldrich (St. Louis, MO). Other chemicals were purchased from Sigma-Aldrich or Affymetrix (Cleveland, OH).

### Fabrication of PDMS channels

Experiments were performed in microfluidic channels and constructed using the toner transfer method as described earlier [22]. The channel pattern depicted in **Figure 1** was drawn in Adobe Photoshop at 1200 dpi resolution with each square pixel ∼21 µm on a side. Channels were 168 x 7581 µm. The pattern was printed onto Toner Transfer Paper (Pulsar, Crawford FL) using a Samsung Laserjet printer (model ML-3471ND) at 1200 dpi resolution. Toner was then transferred using pressure and heat to a brass substrate cut from shim stock (0.032 inch thick, Amazon). Brass etching was performed with a 20% solution (% w/v) ammonium persulfate (APS, Sigma). Regions protected by toner are not etched. We etched to depths of 20-40 µm using the total substrate weight to monitor progress. The depth was occasionally measured experimentally using the narrow depth of focus for the TIRF objective as described previously [23]. When making the PDMS substrate, Sylgard 184 base and curing agent were mixed extensively in a 10:1 ratio, degassed in a vacuum pump, and poured over the brass mold in a Petri dish. A microchip socket was put on the top of the mold before pouring with each end of the channel in the brass mold contacted by one pin of the socket. After curing for 60 hours at 21°C, the socket was removed and the PDMS substrate with channels was carefully peeled off the brass mold. (Although curing at higher temperature speeds up the process, we found that PDMS chips obtained from slow curing made the better seal on NC coated glass.) The socket pin left an access port to the channel through the PDMS substrate.

**Figure 1.**
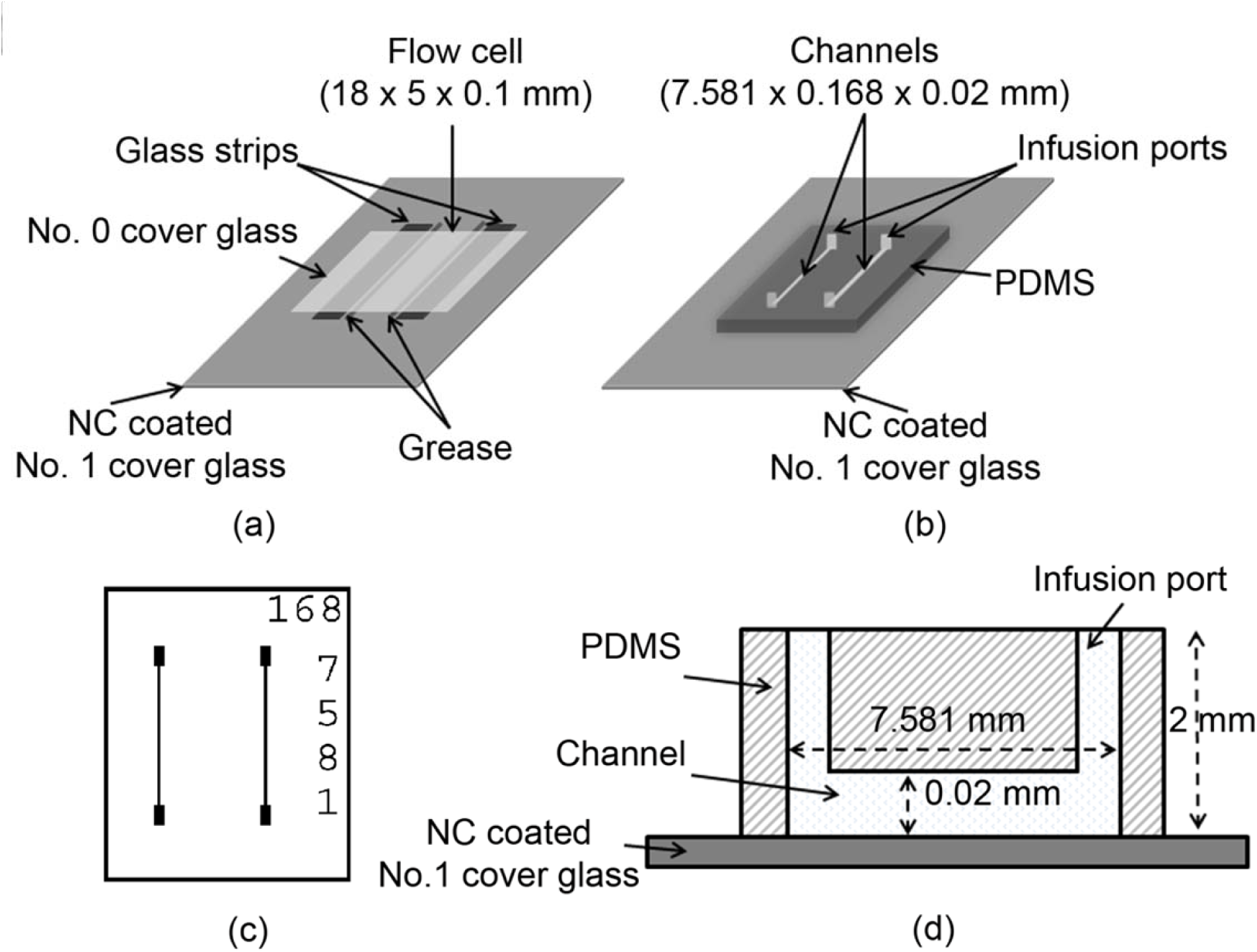
The diagrams of flow cells. (a) The diagram of standard glass flow cell. (b) The diagram of PDMS flow cell. (c) The PDMS channel pattern. (d) The sketch of PDMS flow cell. There are two channels in each PDMS chip. The channel dimension is 168 µm × 7581 µm. Channel depth is 20 µm.

### Protein preparations

Rabbit skeletal muscle proteins were prepared from back and leg muscles of male New Zealand white rabbits. Myosin was prepared by the method of Tonomura et al. [24] and sHMM obtained by chymotryptic digestion of myosin [25]. G-actin was obtained from rabbit skeletal muscle acetone powder by using the method described by Pardee and Spudich [26] then stored immediately under argon gas in liquid nitrogen. Before actin was used, the frozen G-actin was thawed and spun at 160,000×g for 90 min to remove denatured actin. Rhodamine labeling of actin filaments was performed with rhodamine-phalloidin and actin in a 1.2:1 molar ratio as described [1]. Biotin-XX-phalloidin plus rhodamine-phalloidin labeling of actin filaments was performed as described previously [27]. Quantum dot (Qdot) streptavidin conjugation to the biotin-XX-phalloidin labeled actin was done in the motility flow cell.

### sHMM in vitro motility

In vitro motility was performed as described previously at 21°C [27]. In vitro motility of Qdot+rhodamine-phalloidin labeled actin was observed with through-the-objective total internal reflection fluorescence (TIRF) [28] on an Olympus IX71 inverted microscope using a 150X, 1.45 NA objective. Images were acquired with an Andor EMCCD camera (iXon3 897 with 16×16 μm pixels and 16 bit dynamic range) using the software supplied by the manufacturer (SOLIS). Time intervals between two frames were 1 s. The actin sliding velocities were analyzed manually using the ImageJ (NIH, USA) plugin MtrackJ [29].

### Construction of flow cells

The glass flow cells were constructed as described [1]. Briefly, two strips of No. 0 thickness cover glass (Thomas Scientific, Swedesboro, NJ) were placed as tracks on a No. 1 thickness glass coverslip (Warner Instruments, Hamden, CT) with NC coated side up. High vacuum grease (Dow Corning, Midland, MI) was placed on the side of strips as sealing. A No. 0 thickness cover glass was placed on the top of the tracks and pressed gently to create a tight seal. The PDMS flow cells were constructed by pressing a PDMS chip (channels side down) onto a NC coated glass coverslip forming a PDMS/glass hybrid flow cell with reversible adhesion of PDMS to glass.

### Coating PDMS surface with nitrocellulose

The PDMS channels were coated with NC the same way as glass was coated [1]. Briefly, two drops of 1% solution of NC in amyl acetate was spread over the surface of water in a 8 cm glass petri dish. The PDMS chip placed on top of the NC film was lifted off with an intact NC coating using plastic wrap stuck to the back of the chip. The NC coated PDMS chip was placed on a paper towel film side up for drying.

### Coating PDMS surface with DDM

The PDMS chips were coated with DDM as described previously with modifications [21]. Both dynamic coating and pre-coating was tested. For the dynamic coating method 1, sHMM solution with 1% DDM was infused into flow cell at the beginning of the motility assay then the standard assay was performed. In the dynamic coating method 2, 1% DDM was in the flow cell all of the time during motility measurements. In the pre-coating process, the PDMS chip was first incubated in 25 mM KCl, 25 mM imidazole pH7.4, 5 mM MgCl_2_, 0.1 mM EGTA, 10 mM DTT, 0.1 mM PMSF (C-buffer) with 0.1% DDM for 5 min then washed with C-buffer. The PDMS chip was then immersed in rapidly stirred C-buffer for 1 hour with the buffer exchanged every 5 min. The PDMS chip was dried with nitrogen and used right away.

### Solution-phase oxidation of PDMS

The PDMS chips were oxidized with H_2_O_2_ as described previously with modifications [20]. The PDMS chips were incubated in mixture of H_2_O/H_2_O_2_/HCl (in a volume ratio of 5:1:1) for 40 min. The PDMS chips were used immediately after purging with deionized water and dried with nitrogen.

### Air plasma pre-treatment

The pristine PDMS chips were used immediately after being air plasma treated for 60s (Harrick Plasma Cleaner PDC-32G).

### 20 day aged PDMS

The pristine PDMS chips were placed in a chamber under ambient condition for 20 days before being used.

### LBL deposition

LBL deposition was performed as described [19]. Briefly, after the PDMS chips were air plasma treated for 90 s, they were immediately filled with a 2% PVA–5% (wt%) Glycerol (PVA/Glycerol) aqueous solution and then placed at 21°C for 20 min. The channels in PDMS chips were emptied by using a vacuum pump and then placed in an oven at 60°C for 2 h. The above steps were repeated once to get the second layer coating. The final coating was immobilized in an oven at 100°C for 20 min.

### PDMS extraction

Extracted PDMS chips were obtained by using the method of Lee et al. [30] with subtle modifications. PDMS chips (1.8 cm × 1.5 cm × 0.2 cm) were extracted in triethylamine for 1 day, deswelled in ethyl acetate for 1 day and then acetone for 2 days (100 mL of each solvent at 21°C; solvent was changed once), and dried in an oven at 90°C for 3 days. PDMS chips lost 6.9% mass after extraction.

### Baked PDMS

For the comparison with extracted PDMS, some pristine PDMS chips without extraction were also baked in the oven together with extracted PDMS chips for 3 days.

## RESULTS AND DISCUSSION

In the current work in vitro motility in PDMS/glass flow cells was evaluated for various treatments of the PDMS. The main findings are summarized in **Table 1**. The sliding velocity of actin filaments in PDMS flow cells with each kind of PDMS modification was obtained by averaging the sliding velocities of actin filaments in two or three independent flow cells. The sliding velocity of actin filaments in one flow cell was measured by averaging the speeds of 20 moving filaments. Each actin filament was tracked for 15 s. Error bars show the standard deviation. When counting moving filaments, only filaments in the field of view in the first frame were considered.

**Table 1.** Velocities of actin filaments sliding over sHMM in different flow cells

| Flow cell material | Surface modification method | Acting sliding velocity ($\mu\text{m/s}$) | Percentage of moving actin (%) | Actin label |
| --- | --- | --- | --- | --- |
| Glass | Coated with NC [1] | $1.91\pm0.09$ | 85-91 | Rh-ph |
| PDMS | 20 day aged | $1.81\pm0.02$ | 70-77 | Rh-ph |
| PDMS | Extracted [30] | $1.38\pm0.13$ | 71-81 | Rh-ph |
| PDMS | DDM pre-coating [21] | $1.57\pm0.09$ | 67-71 | Rh-ph |
| PDMS | Solution-phased oxidation [20] | $1.37\pm0.12$ | 65-77 | Rh-ph |
| PDMS | LBL [19] | $1.40\pm0.02$ | 60-77 | Rh-ph |
| PDMS | Coated with NC | $1.27\pm0.19$ | 60-67 | Rh-ph |
| PDMS | 60s air plasma pre-treatment [30] | $1.16\pm0.04$ | 57-60 | Rh-ph |
| PDMS | Baked | $1.03\pm0.08$ | 57-65 | Rh-ph |
| PDMS | DDM dynamic-coating 1 [21] | $1.21\pm0.02$ | 41-51 | Rh-ph |
| PDMS | None | $0.80\pm0.61$ | 9-51 | Rh-ph |
| PDMS | DDM dynamic-coating 2 [21] | $0.62\pm0.10$ | <15 | Rh-ph |
| PDMS | 20 day aged | $1.42\pm0.02$ | 14-32 | Qdot |
| PDMS | Extracted [30] | $1.31\pm0.17$ | 56-57 | Qdot |
| Glass | Coated with NC | $1.90\pm0.06$ | 61-67 | Qdot |

Glass flow cells were used as control. In control flow cells, the average velocity of Rh-ph labeled actin sliding over sHMM at 100 µg/ml bulk sHMM concentration was 1.91 µm/s, and 85-91% of the total actin filaments were able to move. In the pristine PDMS flow cells, the average actin sliding velocity was reduced to 0.80 µm/s, and only 9-51% actin filaments moved. The poor motility in pristine PDMS flow cells suggested that the PDMS modification is necessary for making in vitro motility PDMS devices practical. In all the PDMS modifications tested, 20 day aged PDMS gave the best overall result. The actin sliding velocity was 1.81 µm/s and 70-77% actin filaments moved. The percent of moving actin filaments was comparable to that in a conventional motility assay using a glass flow cell where 80-95% moving actin filaments is typical [31; 32; 33]. The sHMM in vitro motility characteristics measured in the conventional glass cell were recovered when using the PDMS flow cell microfluidic when the PDMS component underwent aging for 20 days making the PDMS a practical substitute for glass.

Extracted PDMS had the highest percentage of moving actin filaments (71-81%) in the test. It also had relatively lower actin sliding velocity (1.38 µm/s). This low velocity might be caused by the low surface density of myosin head, which is due to depletion of the bulk myosin concentration from protein adsorption to PDMS. As a low duty cycle myosin, the velocity of actin sliding over sHMM decreases with decreasing surface density when the surface density is below threshold [34; 35; 36]. In the NC coated glass surface, the velocity starts to decrease once bulk sHMM concentration is below 100 µg/ml [27; 36].

Surface area and hydrophobicity are likely important factors impacting how PDMS affects motility. The PDMS surface appears irregular in the microscope under high magnification compared to cover glass implying it presents higher surface area for protein adsorption. The PDMS flow cells have much smaller dimension than the standard glass flow cell does, which makes PDMS flow cells have much higher surface area to volume ratio. Protein affinity for the surface depends on surface charge assessed macroscopically with the contact angle for water. Experiment shows that the difference in contact angle of water on PDMS surface between pristine PDMS and 20 day aged PDMS is negligible [16]. As a result, the big difference in motility between them is more likely related to the larger surface area to volume ratio and implying leaching from the PDMS could be a factor. It has been shown previously that at least 5% (w/w) of the PDMS bulk remains uncrosslinked after extensive curing [16]. Uncured PDMS oligomers were reduced but still detectable in microchannel media after incomplete extraction of PDMS (PDMS chip lost 4% mass through overnight Soxhlet extraction in ethanol in that experiment) [37]. PDMS curing is a time dependent process. 20 day aging probably reduces the amount of uncured oligomer in PDMS chips. Consequently, the amount of uncured oligomers entering the medium in the flow cells built with 20 day aged PDMS is also reduced. We hypothesize that the contamination from the leaching of uncured PDMS oligomers into the sample media is one of the major reasons motility is compromised. The good recovery in the percentage of moving actin filaments using extracted PDMS flow cells (**Table 1**) supported the hypothesis. Neutralizing uncured oligomers can also be achieved by Soxhlet extraction and extended oven baking, besides serial extraction in organic solvents [30; 38; 39].

Both air plasma oxidation and solution oxidation improved motility in PDMS flow cells (**Table 1**). Repeating units of –O–Si(CH_3_)_2_– is the cause of hydrophobicity of PDMS surface [9]. Oxidizing PDMS introduces silanol (Si-OH) groups and removes methyl (Si-CH_3_) groups, thus lowering surface hydrophobicity [9; 40; 41; 42]. Contact angle of water on PDMS surface can be successfully reduced from ∼98° to ∼42° after treatment with an air plasma for 60 s [30]. Better reductions on contact angle were achieved using extracted PDMS or oxygen plasma [30; 43; 44; 45; 46; 47; 48; 49]. The silica-like surface layer formed during oxidation was shown to reduce O_2_ permeability of the chamber preventing motor protein oxidation [14] but also might slow down the uncured PDMS oligomers entering solution.

To build a myosin driven microfluidic based assay, 20 day aging is a simple and effective PDMS surface modification technique requiring no instrumentation. In a 20 day aged PDMS channel which has a volume of 0.03µl, the actin sliding velocity was 1.81 µm/s and fraction of moving actin filaments was 70-77%. A quite recent work applied a 40 min air plasma pre-treatment of PDMS in vitro motility flow cell [14]. It achieved similar results with a much larger microchannel (∼1.2 µl), more specialized instruments (i.e., spin coater and air plasma cleaner), and less wait time. In that work, the actin sliding velocity was ∼1.6 µm/s in a PDMS/glass microchannel. In a PDMS/glass flow cell in which the PDMS microchannel was replaced by a PDMS slab, the actin sliding velocity was ∼1.7 µm/s and the percentage of moving actin filament was ∼70%. Poly(methyl methacrylate) PMMA coated with trimethylchlorosilane (TMCS) is a reasonable alternative choice for construction of microfluidic devices [50], if specialized instruments (i.e., spin coater, electron beam lithography system and oxygen plasma cleaner) are accessible.

Qdot fluorescent tags have great advantages in applications involving super resolution microscopy because they are much brighter and much less susceptible to photobleaching than organic dyes and fluorescent proteins [51; 52]. We introduced the Qdot to the in vitro motility assay and used super resolution microscopy for rapid, quantitative, and inexpensive step-size measurement in low duty cycle muscle myosins implicated in inheritable muscle diseases [3; 27]. In the flow cells built with 20 day aged PDMS or extracted PDMS, the velocity and percentage of moving actin filaments in motility using Qdot labeled actin declined compared to that using rhodamine labeled actin. Further investigations on how to modify PDMS to improve in vitro motility assay with Qdot labeled actin are needed.

## ETHICS STATEMENT

This study conforms to the Guide for the Care and Use of Laboratory Animals published by the US National Institutes of Health (NIH Publication no. 85–23, revised 2011). All protocols were approved by the Institutional Animal Care and Use Committee at Mayo Clinic Rochester (protocol A56513, effective September 2, 2016).

## RESEARCH DATA

Data used in this study is provided in summary form in **Table 1**. Public domain ImageJ software was used for analysis as noted in the text. Representative raw in vitro motility and Qdot assay data were deposited at Zenodo (search: cc01bdb351e7ba0a8562849935a1e54e). Metadata accompanies the raw data identifying samples and parameters for image capture.

## AUTHORS’ CONTRIBUTIONS

YW conceived and designed experiments, performed experiments, analyzed data, and co-wrote the paper with TPB. TPB conceived and designed experiments and co-wrote the paper with YW. Both authors gave final approval for publication.

## ACKNOWLEDGMENT

This work was supported by NIH grants R01AR049277 and R01HL095572 and by the Mayo Foundation

